# Fine-scale genetic structure and rare variant frequencies

**DOI:** 10.1101/2024.02.02.578687

**Authors:** Laurence Gagnon, Claudia Moreau, Catherine Laprise, Simon L. Girard

## Abstract

In response to the current challenge in genetic studies to make new associations, we advocate for a shift toward leveraging population fine-scale structure. Our exploration brings to light distinct fine-structure within populations having undergone a founder effect, challenging the prior perception of homogeneity. This underscores that smaller, but well-defined cohorts, demonstrate an important increase in rare variant frequencies, offering a promising avenue for new genetic variants’ discovery.

## Text

Common variants are the primary source of variation identified by genetic association studies. However, despite numerous association analyses conducted in the last years, a significant proportion of the genetic predisposition for many diseases still remains unknown. To address this issue, it is crucial to study rare variants, which often have more significant phenotypic effects, to better understand this “missing heritability”. Nevertheless, associations with rare variants present a reduction in statistical power due to the scarcity of individuals carrying these alleles^1^. Thus, given their unique structure, populations that had undergone a founder effect (PFE) have the potential to more readily reveal new genetic associations of rare variants that have potential implications for human health^2–7^, and may help to reduce the “missing heritability”^8^. Therefore, our study aims to properly describe the fine-scale genetic structure of a population to refine analysis models for genetic associations.

We characterized the genetic structure of four PFE alongside three reference groups from the 1000 Genomes Project. These populations are the Himba of Namibia, the Hutterites of South Dakota in the US, the population of the Quebec’ province in Canada and the Ashkenazi Jews of Europe and across the World (Methods Table 1). A Uniform Manifold Approximation and Projection (UMAP) analysis was conducted to analyze their genetic structure (Fig.1A). The Himba and Hutterites form two tightly packed clusters, whereas the Ashkenazi Jews and Quebec exhibit a more dispersed pattern and are even interconnected. For Hutterites and Himba, they can be studied as a whole as they do not subdivide into clusters. Indeed, Hutterites are known to practice endogamy and live in community and Himba individuals were documented to practice polygyny and live in a pastoralist way^9,10^. This way of living promotes very close links between individuals and could explain the absence of fine-structure. This is also evident in the proportion of pairs sharing an identity-by-descent (IBD) segment across the genome, which reaches the highest level among the Himba and Hutterites compared to the other PFE. In contrast, a fine-structure is observed within the Ashkenazi Jews and the Quebec population so that they can be subdivided into clusters (Fig 1B). The fine-structure in these populations is also reflected in the patterns of sharing of IBD segments; indeed, some of the clusters exhibit increased IBD sharing across the genome (Extended Table 1). The proportion of pairs sharing an IBD segment through the genome within these clusters reaches an average threshold comparable to the one observed among Himba and Hutterites (Extended Table 1). The Ashkenazi Jews and Quebec went through unique histories of migration, isolation and population expansion. The Ashkenazi Jews-1 cluster represents the Ashkenazi Jewish ancestry, i.e. individuals for whom all four grand-parents were of Ashkenazi Jewish ancestry (Extended Figure 2A). In contrast, the Ashkenazi Jews-2 cluster appears to represent a more admixed ancestry, due to its connection to the European reference group and Quebec (Fig. 1B). This cluster likely contains individuals with only 1 to 3 of their grand-parents with Ashkenazi Jewish ancestry^6^. Moreover, the analysis of the genetic relatedness among and between clusters reveals that the Ashkenazi Jews clusters are distinct and exhibit greater relatedness within than between clusters (Extended Figure 3A). As for Quebec, clusters can be associated with specific ethnocultural groups. Specifically, the Quebec-2 and Quebec-3 represent the Saguenay-Lac-Saint-Jean and the Acadians of Gaspe, respectively (Extended Figure 2B). These two groups are known for having a genetic structure which distinguishes them from the broader Quebec population that can be associated with Quebec-1 cluster^11^. This cluster is related to the initial founder effect in Quebec. This is evident in the lower genetic relatedness observed within the Quebec-1 cluster compared to all the others (Extended Figure 3B). Undoubtedly, populations like the Ashkenazi Jews and Quebec cannot be treated as single entities due to the presence of fine-structure, even if they were initially perceived as “homogenous populations”.

**Table 1.** Frequency of disease-causing variants known to be associated with a specific population. Only the main population and the associated cluster are shown on the table.

| Population | MAF of the European reference group | MAF of the population | Number of individuals |
| --- | --- | --- | --- |
| Familial dysautonomia (chr9:108899816:A:G) (ClinVar: 6085) |  |  |  |
| Ashkenazi Jews | 0.0000 | 0.0178 | 2052 |
| Ashkenazi Jews-1 | 0.0000 | 0.0211 | 1729 |
| Spastic ataxia of Charlevoix-Saguenay (chr13:23335031:TA:T) (ClinVar: 5512) |  |  |  |
| Quebec | 0.0000 | 0.0080 | 941 |
| Quebec-2 | 0.0000 | 0.0323 | 233 |
| Usher syndrome type I (chr11:17531431:C:T) (ClinVar: 5143) |  |  |  |
| Quebec | 0.0000 | 0.0048 | 941 |
| Quebec-3 | 0.0000 | 0.0385 | 52 |

**Figure 1.**
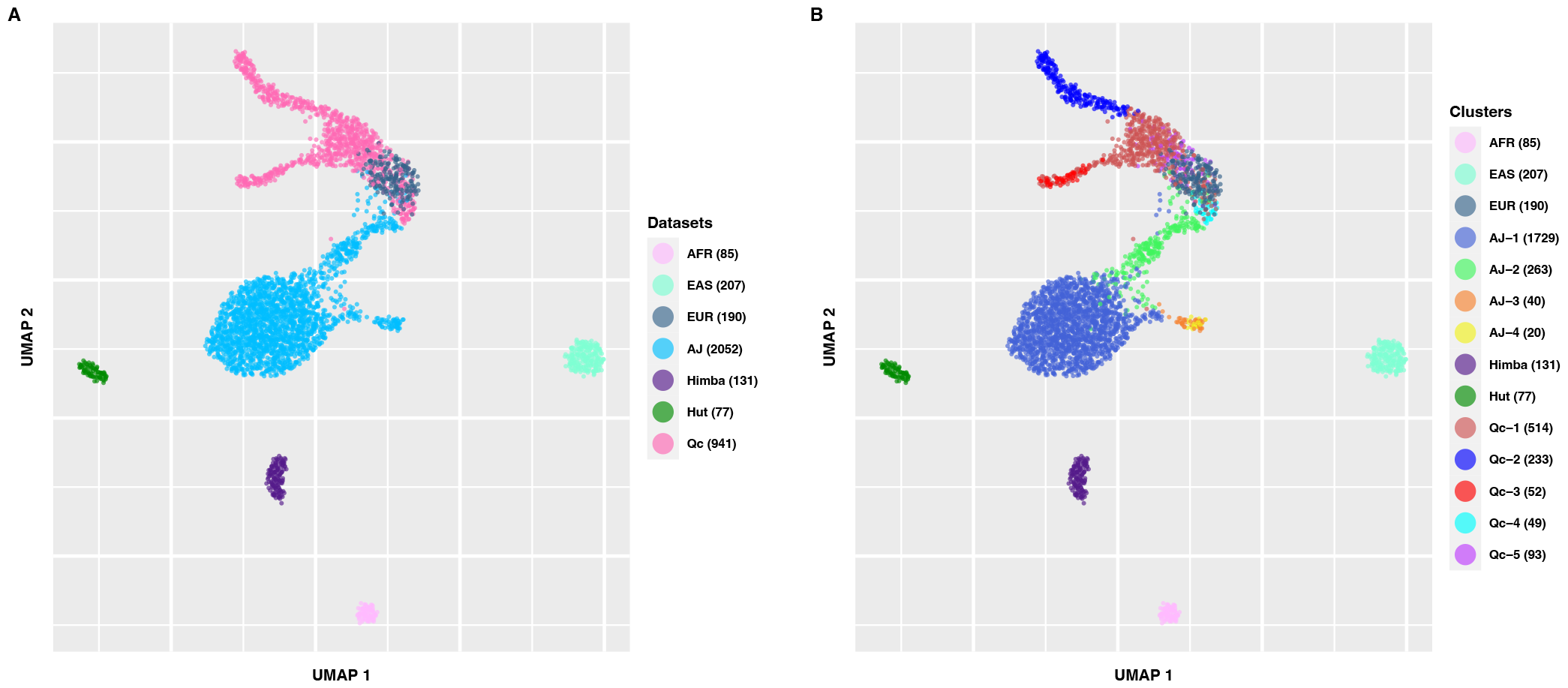
UMAP of the first 8 principal components of the merged dataset. Colored according to the origin of the population (A) and the clusters (on Extended Figure 1) (B). AFR African from the 1000 Genomes Project, EAS East Asian from the 1000 Genomes Project, EUR European from the 1000 Genomes Project, AJ Ashkenazi Jews, Hut Hutterites, Qc Quebec.

To assess the impact of clustering on genetic association, we conducted an analysis on known founder disease variants. These variants were selected for their association within a specific group. The analysis of the frequency of these variants in the associated group (Quebec-2 for Saguenay-Lac-St-Jean, Quebec-3 for Acadians of Gaspe, or Ashkenazi Jews-1 for Ashkenazi Jews) (Extended Figure 2), revealed higher allele frequencies of the variant in the specific cluster compared with the other clusters (Table 1). They were also nearly absent from the European reference group. Remarkably, within the Quebec population, the variants associated with spastic ataxia of Charlevoix-Saguenay and Usher syndrome type I exhibit a 4 and 8 fold increase, respectively, when comparing the specific cluster and the whole population. Notably, this was calculated with a much smaller sample size of 4 and 18 fold, respectively. This trend is also observable while investigating the variant of Familial dysautonomia in the Ashkenazi Jews and other diseases associated with a specific population (Table 1 and Extended Table 2). So, it is crucial to consider these clusters not only in PFE, but also in outbred populations since the presence of clusters in more diverse or admixed populations could even have a more striking effect on genetic associations^12,13^. This approach could result in the need for much smaller cohorts consisting of individuals with well-known fine-scale genetic structure, offering cost-effectiveness and increased statistical power.

In conclusion, we suggest a novel approach, parallel to the already existing strategies, that uses the fine-scale genetic structure of a population to refine analysis models for genetic associations. This cost-effectiveness method would help to enhance the value of existing large cohorts and to develop new analytical methods. We believe that cohorts composed of fewer individuals with a common genetic background would help in discovering new rare genetic associations, as they would be easier to find given their increased frequency. Investigating more prevalent diseases within targeted populations has the potential to generate positive impacts on public health at the community level and on the discovery of new genes that could be new therapeutic targets.

## Data availability

The Quebec cohort genotypes are available upon request to BALSAC at https://balsac.uqac.ca/^14^. The Ashkenazi Jews cohort data may be available via dbGaP study accession number phs000448.v1.p1, the Himba cohort data may be available via dbGaP study accession number phs001995.v1.p1 and the Hutterites cohort data may be available via dbGaP study accession number phs001033.v1.p1.

## Code availability

The code used for this study can be found in the following GitHub repository: https://github.com/laugag17/world_pop_with_founder_effect.

## Acknowledgements

This work was supported by funding the Canada Research Chair in Genetics and Genealogy in the SLG lab. It was also made possible by the Digital Research Alliance of Canada which provided access to storage and computing resources. We are extremely grateful to all participants of this research. We would like to thank Hélène Vézina, Damian Labuda and their team for the Quebec Regional Reference Sample cohort constitution. LG received funding from the Fonds de recherche du Québec - Santé and the Canadian Institutes of Health Research. CL is the director of the Centre intersectoriel en santé durable (http://www.uqac.ca/santedurable), the chairholder of the Canada Research Chair in the Genomics of asthma and allergic diseases (http://www.chairs.gc.ca) and co-holder of the Chaire en santé durable du Québec (http://www.chairesantedurable.ca).

## Author contributions

All authors acquired data and approved the final version of the manuscript. LG and CM played an important role in interpreting the results. LG, CM and SLG conceived and designed the study and drafted the manuscript. CL, designed, built and manages the Saguenay-Lac-Saint-Jean cohort, obtaining funding for genealogical constructs and genomic data acquisition. CL also revised the manuscript.

## Subjects and methods

This study was approved by the University of Quebec in Chicoutimi (UQAC) ethics board.

### Cohorts

The data consist of 4 different cohorts of populations that had undergone a founder effect (PFE). We gained access to data from Quebec, Ashkenazi Jews, Himba and Hutterites (Methods Table 1). We also used the data from the 1000 Genomes Project as reference groups from Africa (Mende (MSL)), Europe (British, Northern and Western Europe (GBR and CEU)) and East Asia (Han Chinese and Japanese (CHB and JPT)). Thus, this led to a final sample size of 3,683 individuals.

**Methods Table 1.** Populations and datasets.

| **Cohort** | **Sample size** | **Populations included** | **Genetic data** | **Sources** |
| --- | --- | --- | --- | --- |
| Quebec | 941 | French Canadian; English-speaking United Empire Loyalists and Acadians of the Gaspé Peninsula. | Illumina Omni Express or Illumina Omni 2.5 chips. | Quebec Regional Reference Sample<sup>11</sup> and unaffected individuals from the Saguenay-Lac-St-Jean asthma familial cohort<sup>15</sup>. |
| Ashkenazi Jews | 2,052 | Ashkenazi Jews. | Illumina HumanOmni1-Quad arrays. | dbGaP study accession number phs000448.v1.p1. |
| Himba | 131 | Himba. | Illumina MEGAex and H3Africa arrays. | dbGaP study accession number phs001995.v1.p1. |
| Hutterites | 77 | Hutterites. | SNP genotypes derived from sequence data. | dbGaP study accession number phs000185.v8.p1. |
| African reference group | 85 | Mende of Sierra Leone (MSL). | SNP genotypes derived from sequence data. | 1000 Genomes Project<sup>16</sup>. |
| European reference group | 190 | British (GBR), Northern Europe and Western Europe (CEU). | SNP genotypes derived from sequence data. | 1000 Genomes Project<sup>16</sup>. |
| East Asian reference group | 207 | Han Chinese (CHB) and Japanese (JPT). | SNP genotypes derived from sequence data. | 1000 Genomes Project<sup>16</sup>. |

### Genotyping data and cleaning

Each dataset underwent cleaning using PLINK software v1.9, ensuring individuals with at least 95% genotypes among all SNPs were retained^17^. At the SNP level, we retained SNPs with at least 95% genotypes among all individuals, located on the autosomes and in Hardy–Weinberg equilibrium p > 0.001 (calculated on each whole cohort). A Principal Component Analysis (PCA) was done on each individual dataset using SNPs with a minor allele frequency (MAF) of at least 5%, and after pruning to remove SNPs in linkage disequilibrium.

Subsequently, all datasets were merged (lifting over to hg19 for the Ashkenazi Jews) to retain only common SNPs. After the merge, individuals with less than 95% genotypes among all SNPs and SNPs with less than 95% genotypes across all individuals were once again filtered out. The final dataset comprises 199,238 SNPs and 4,259 individuals. Related individuals (PLINK pihat >= 0.25) were filtered out, resulting in a final sample size of 3,683 subjects. This unusually high threshold was applied to retain two populations with high relatedness (Extended Table 3). Hutterites are indeed recognized for practicing endogamy and communal living, while Himba individuals have a pastoralist lifestyle and practice polygyny^9,10^. The Extended Figure 4 demonstrates the low impact of different genetic relatedness thresholds on the pairwise sum of identity-by-descent (IBD) segments length and number. Finally, a PCA was performed on SNPs with a MAF of at least 5% and after pruning to remove SNPs in linkage disequilibrium (73,624 SNPs left).

The merged dataset was imputed on TOPMed imputation server, using the reference panel topmed-r2 after lifting over to hg38^18^. MAF of known disease-causing variants were computed using PLINK software v1.9. It’s important to note that using imputed data may result in a loss of rare variants or in the underestimation of the real frequency of these variants in a PFE. The founder variants (listed in Table 1 and Extended Table 2) were selected because of their association with specific populations. However, limited literature is available concerning specific variants in the Acadians of Gaspe (Quebec-3) which could explain why we found only one variant associated of interest in this cluster.

### Statistical analysis

A Uniform Manifold Approximation and Projection (UMAP) was performed on the first 8 principal components of the PCA to capture as much variance as possible. Additional UMAPs were completed on each individual dataset, using the first 5, 4, 5, 8 principal components for the Ashkenazi Jews, Quebec, Himba and Hutterites, respectively (Extended Figure 1). The UMAPs were realized with the R package “umap” v0.9.2.0^19^. The n neighbors variable was set to the maximal value (number of individuals in the dataset) (Methods Table 1). The min distance value was set to 0.9 for the UMAP on the merged dataset to promote dots splitting and ensure good visualization; while it was set to 0.01 for the UMAPs on individual datasets of populations to better capture the structure and promote clustering^20^.

The DBScan method was employed to generate clusters from UMAP of each population. This method was chosen for its ability in density-based clustering, allowing the capture of clusters with various shapes, including non-convex shapes^21^. The “dbscan” R library v1.1-11 was used for clustering with the minPts parameter set to 4, and the epsilon value adjusted according to each individual population. The Himba and Hutterites only displayed a single cluster each, while Ashkenazi Jews and Quebec exhibited 4 and 5 clusters, respectively (Extended Figure 1).

### Analyses of IBD segments

The assessment of pairwise IBD segments was performed using refinedIBD software v17Jan20 on phased genotypes, which was done using Beagle software version 18May20.d20^22^. The identified segments were then merged with merge-ibd-segments.17Jan20.102.jar. This software was selected for its robustness and precision in detecting IBD segments^22^. Only segments of 2 cM or more and with a LOD score greater than 3 were retained for further analysis on the level of IBD sharing across the genome.

