## Extended information for "Fine-scale genetic structure and rare variant frequencies"

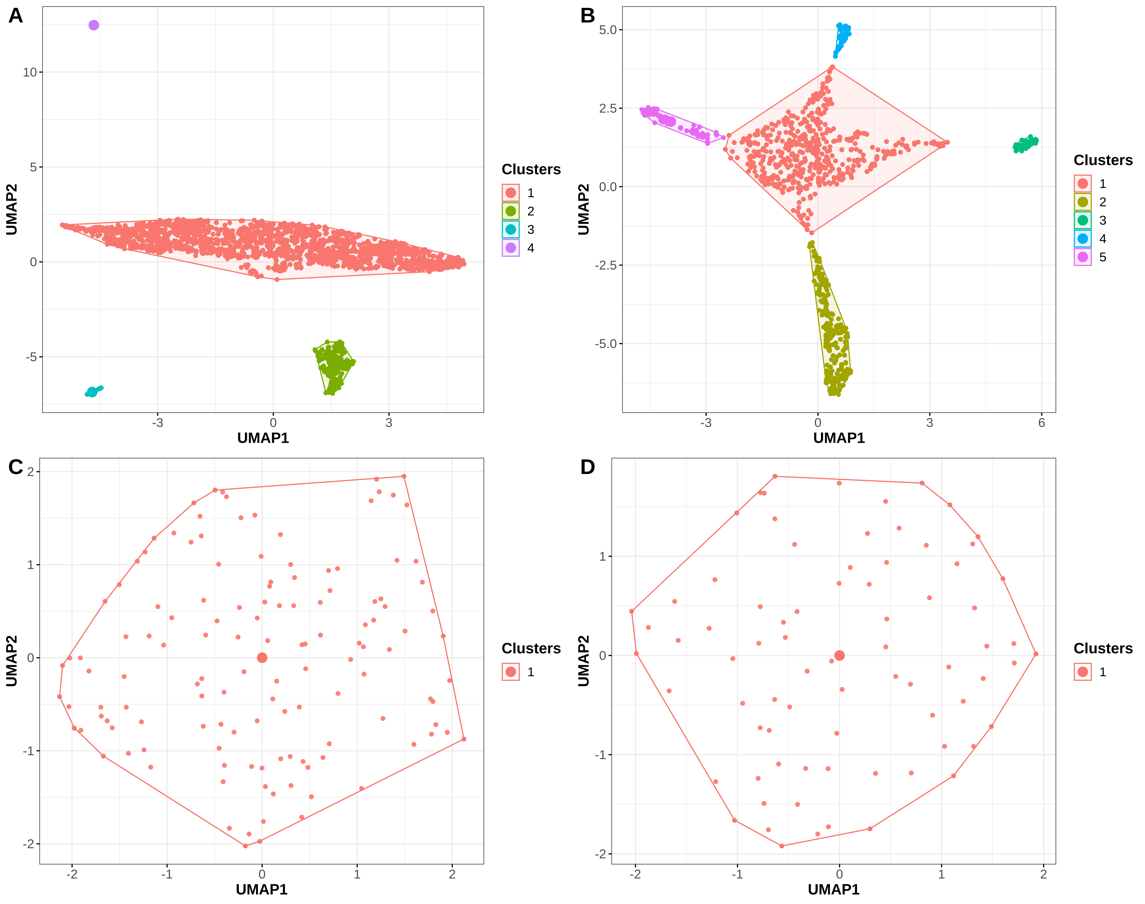

**Extended Figure 1.** UMAP clustering with DBScan for each dataset of PFE.

A. Ashkenazi Jews. B. Quebec. C. Himba. D. Hutterites

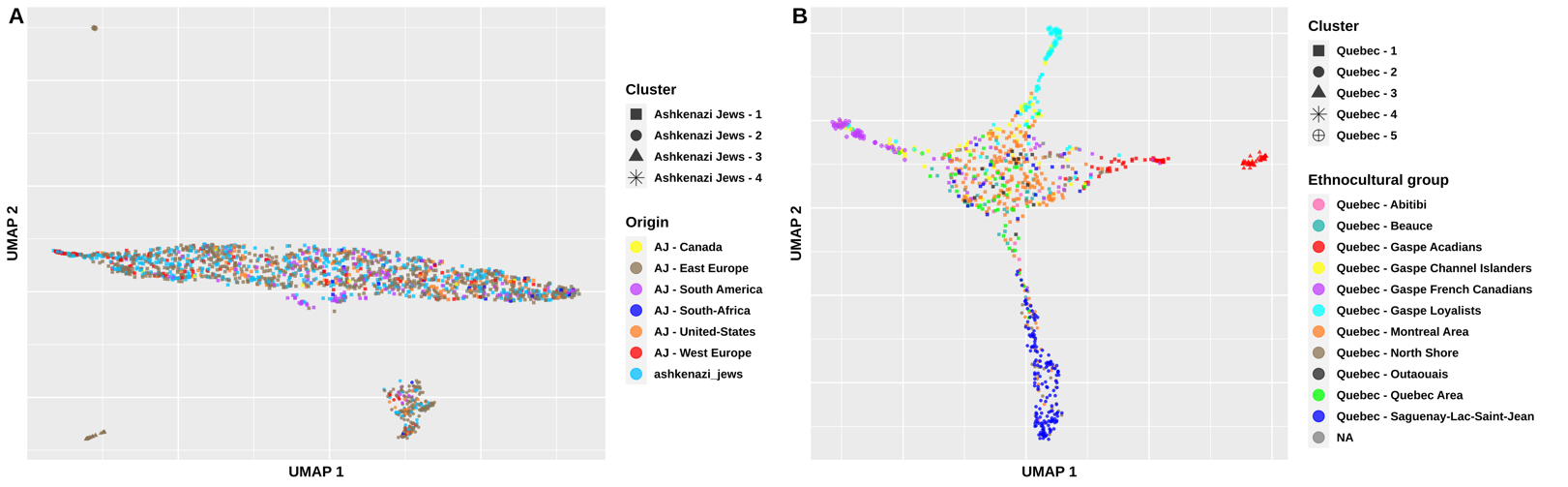

**Extended Figure 2.** UMAP colored according to the origin or ethnocultural group and shaped according to the clustering.

A. Ashkenazi Jews. B. Quebec. For panel A, Ashkenazi Jews have reached these locations already long after the nearly complete mixing of the population in Eastern and Central Europe. Therefore, no genetic differences were expected across these locations^6^.

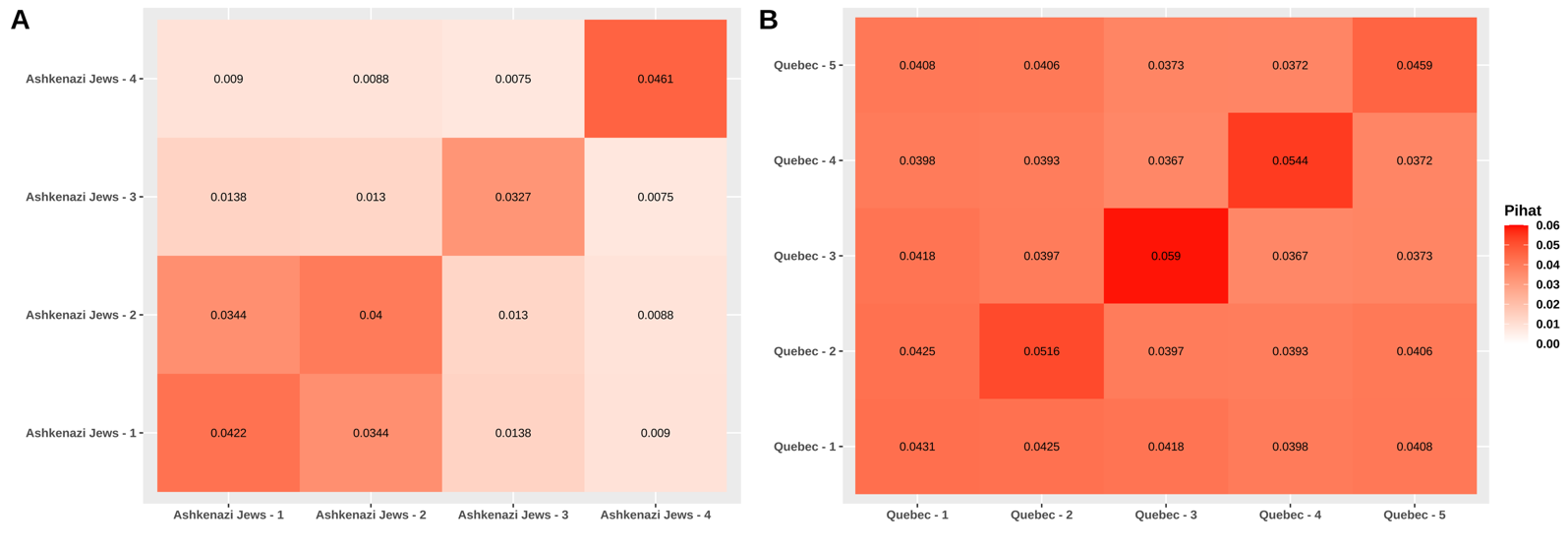

**Extended Figure 3.** Heatmap of the averaged genetic relatedness (PLINK pihat) between and within clusters.

A. Ashkenazi Jews. B. Quebec.

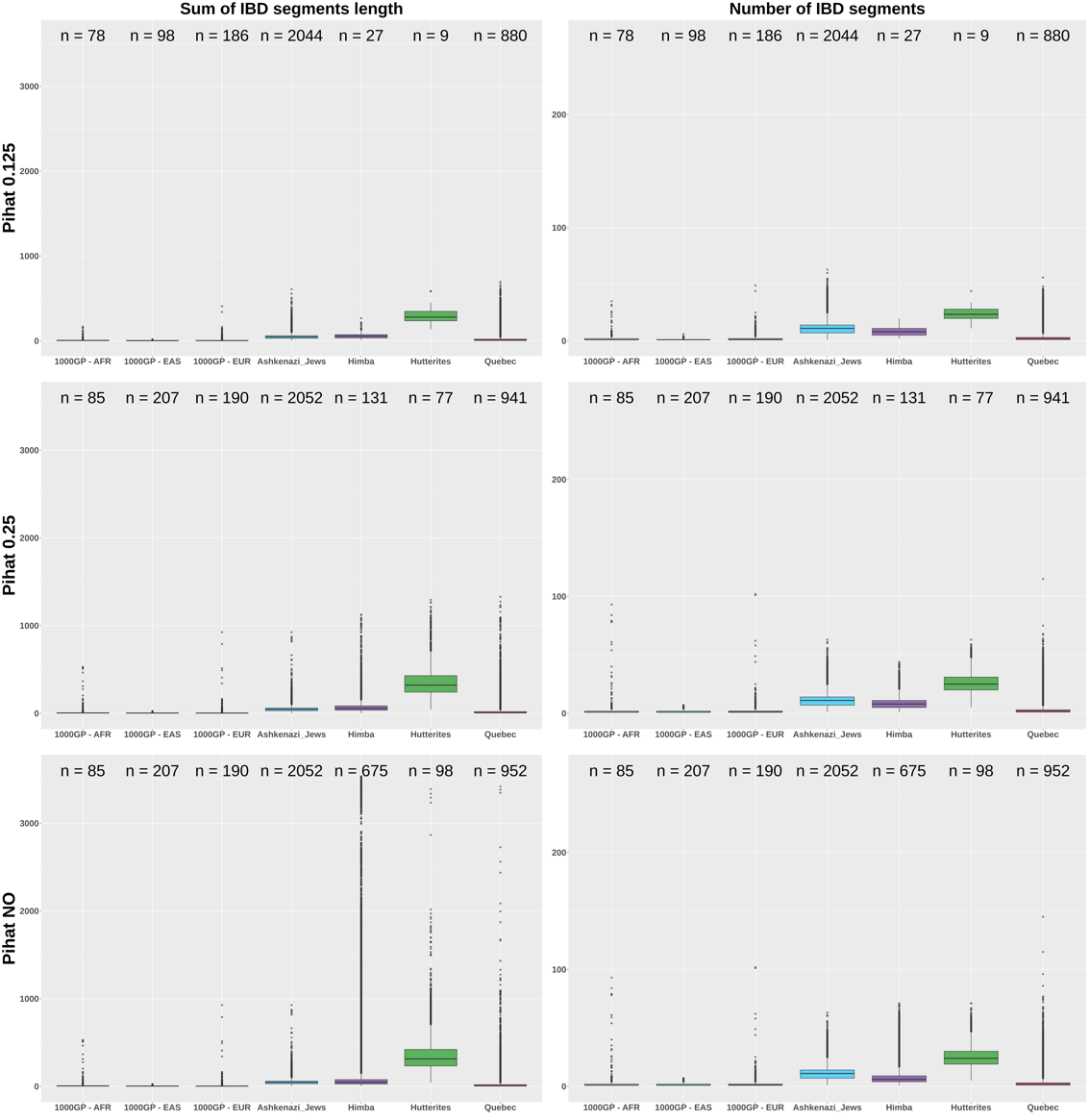

**Extended Figure 4.** Pairwise sum of IBD segments length and number of IBD segments according to different genetic relatedness filters (pihat 0.125, pihat 0.25 and no cleaning).

**Extended Table 1.** Mean, maximum and minimum proportion of pairs sharing an IBD segment through the genome for each PFE, clusters and reference population.

| **Population** | **Mean (%)** | **Maximum (%)** | **Minimum (%)** |
| --- | --- | --- | --- |
| Himba | 2.182 | 3.464 | 0.611 |
| Hutterites | 10.309 | 14.730 | 2.461 |
| Quebec | 0.396 | 0.819 | 0.073 |
| Quebec - 1 | 0.251 | 0.606 | 0.032 |
| Quebec - 2 | 2.130 | 3.630 | 0.359 |
| Quebec - 3 | 4.897 | 10.106 | 1.207 |
| Quebec - 4 | 3.013 | 8.078 | 0.595 |
| Quebec - 5 | 1.653 | 3.296 | 0.234 |
| Ashkenazi Jews | 1.211 | 1.770 | 0.081 |
| Ashkenazi Jews - 1 | 1.524 | 2.196 | 0.105 |
| Ashkenazi Jews - 2 | 0.261 | 0.633 | 0.012 |
| Ashkenazi Jews - 3 | 2.520 | 10.256 | 0.128 |
| Ashkenazi Jews - 4 | 5.986 | 27.895 | 0.526 |
| 1000GP - EUR | 0.094 | 1.470 | 0.006 |
| 1000GP - EAS | 0.048 | 1.351 | 0.005 |
| 1000GP - AFR | 0.171 | 1.261 | 0.028 |

**Extended Table 3.** Number of individuals remaining according to different genetic relatedness filters.

| **1000GP - AFR** | **1000GP - EAS** | **1000GP - EUR** | **Ashkenazi Jews** | **Himba** | **Hutterites** | **Quebec** | **Genetic relatedness** |
| --- | --- | --- | --- | --- | --- | --- | --- |
| 85 | 207 | 190 | 2052 | 675 | 98 | 952 | No filter |
| 85 | 207 | 190 | 2052 | 131 | 77 | 941 | 0.25 |
| 78 | 98 | 186 | 2044 | 27 | 9 | 880 | 0.125 |
